# Allometric scaling of a superposition eye optimises sensitivity and acuity in large and small hawkmoths

**DOI:** 10.1101/2022.02.08.479593

**Authors:** Anna Stöckl, Rebecca Grittner, Gavin Taylor, Christoph Rau, Andrew J Bodey, Almut Kelber, Emily Baird

## Abstract

Animals vary widely in body size across and within species. This has consequences in large and small individuals for the function of organs and body parts. How these scale in relation to body size reveals evolutionary investment strategies, often resulting in trade-offs between functions. Eyes exemplify these trade-offs, as they are limited by their absolute size in two key performance features: sensitivity and spatial acuity. Previous studies of the 3D structure of apposition compound eyes, which are ideal models for allometric studies due to their size polymorphism, revealed that allometric scaling improves both local resolution and visual sensitivity in larger bumblebees (Taylor et al., 2019). Here, we build on the established methods and results to investigate allometric scaling in superposition compound eyes – the second prominent eye type in insects – for the first time. Our research highlights a surprising strategy to cope with the challenge of trading off sensitivity and spatial resolution in small eyes, as we show that the eyes of the hummingbird hawkmoth retain an optimal balance of these performance measures across all body sizes.

## Introduction

Animals of the same species can vary considerably in body size (Blanckenhorn, 2000; Chown & Gaston, 2010; Sibly & Brown, 2007). Such differences have performance consequences for body parts or organs in larger and smaller individuals, particularly when their function depends on absolute rather than relative size (Spence, 2009). A key organ that exemplifies the evolutionary strategies to cope with the behavioural and ecological consequences of body size variation is the eye, because eyes are performance-constrained by their absolute size. Eye size, in turn, is limited by body size, due to the energy and weight constraints associated with carrying large eye structure, particularly in small flying animals (Niven & Laughlin, 2008). Eye size limits two central features of eye functionality: sensitivity and spatial resolution (Land et al., 1997; Snyder, 1977; Snyder et al., 1977; Warrant & McIntyre, 1993). Larger eyes can collect more photons, due to a potentially larger light collecting aperture and focal length, as well as the diameter and length of their photoreceptive units. Higher sensitivity is not just important for seeing well in dim light (Warrant & McIntyre, 1993), but also for discriminating fine contrast changes at higher light intensities (Snyder, 1977; Snyder et al., 1977). In addition, spatial resolution is limited by the number of visual units packed into an eye of a given viewing angle – thus the number of “pixels” that can be resolved across the eyes’ field of view (Land et al., 1997; Snyder, 1977; Snyder et al., 1977; Warrant & McIntyre, 1993). While a small eye could densely pack many visual units with high acuity, the small eye size means that they will have to be narrower than in larger eyes, and thus of lower light sensitivity, and consequently lower contrast resolution (Snyder, 1977; Snyder et al., 1977). This size limit on spatial resolution is exacerbated in eyes with small lenses, such as the compound eyes of insects. Here, the small diameter of facets can set a diffraction-limit to the optical resolution, resulting in blurred visual projections (Snyder, 1979; Stavenga, 2003; Stavenga, 2006; Warrant et al., 2007). Combined with their generally small body size that restricts the absolute eye size (Niven & Laughlin, 2008), these challenges to sensitivity and spatial resolution make insect compound eyes an ideal model to study how eyes scale allometrically for optimal performance in small animals.

One strategy that most insect species use to cope with these challenges is to preserve an eye as large as possible in small individuals, resulting in a negative allometric relationship between eye and body size. This means that smaller individuals have absolutely smaller but relatively larger eyes for their body size), within and across species (bees: (Jander & Jander, 2002; Spaethe, 2003; Streinzer & Spaethe, 2014; Taylor et al., 2019), ants: (Perl & Niven, 2016a; Zollikofer et al., 1995), butterflies: (Merry et al., 2006; Rutowski, 2000), and flies (Currea et al., 2018). Positive allometry between eye and body size is rare (Streinzer et al., 2016). A second trend commonly observed in insects is a negative allometry between facet size and eye size (Currea et al., 2018; Merry et al., 2006; Perl & Niven, 2016a; Taylor et al., 2019; Zollikofer et al., 1995). A relatively larger facet size in smaller individuals can improve visual sensitivity (Land et al., 1997). Larger bumblebees, for example, forage at lower light intensities that smaller ones (Kapustjanskij et al., 2007) and detect smaller point-targets because of an increased sensitivity of individual ommatidia (Spaethe, 2003). These scaling strategies do not always manifest over the entire eye, but can also differ locally (Perl & Niven, 2016b). In bumblebees, for example, larger individuals benefit from optimising spatial acuity in their frontal acute zone, while the overall spatial resolution of the eye remains similar in all individuals (Taylor et al., 2019).

All of these insights into the scaling strategies of insect eyes are based on apposition compound eyes, in which the sensitivity of individual optical units is limited by their facet size. A large proportion of insects, however, especially among the Lepidoptera and Coleoptera (Exner, 1891; Kunze, 1972), possesses a different eye type: superposition compound eyes.

This eye type is typically found in nocturnal insects, though with prominent diurnal exceptions. It provides a highly increased sensitivity compared to apposition eyes (Land et al., 1997; Snyder, 1977; Warrant & McIntyre, 1993), since hundreds of neighbouring facets can focus light onto a single rhabdom, acting as a functional lens with an aperture larger than that of a single facet (Exner, 1891). This increased single-ommatidial photon capture might lead to different selection constraints in the scaling with body size compared to apposition eyes (Meyer-Rochow & Gál, 2004). Moreover, because of the intricate optical arrangements of multiple corneal lenses and crystalline cones that focus light onto a single rhabdom, superposition compound eyes might be less flexible for local modifications, as these could compromise the superposition optics. Thus, revealing the scaling strategies of superposition compound eyes will be an important contribution to understanding the visual constraints of many beetle and moths species – many of which are important diurnal and nocturnal pollinators (Kevan & Baker, 1983; Proctor M, 1996).

To quantify how superposition compound eyes scale with body size, we chose to study an insect model that can directly compared to species with apposition eyes: the hummingbird hawkmoth *Macroglossum stellatarum*. As day-active nectar foragers (Stöckl & Kelber, 2019), these moths are under similar visual constraints as many previously tested hymenopteran and lepidopteran species, and share habitats and host plants with common Eurasian bee and butterfly species. To quantify the allometric scaling of optical and sensory structures of the eyes of large and small hummingbird hawkmoths, we used X-ray micro computed tomography (Bagheri et al., 2019; Baird & Taylor, 2017; Taylor et al., 2019). Even though the eyes of hawkmoths are generally designed for high photon catch, we found a strong negative allometry between eye and body size, and between facet diameter and eye size, resulting in a proportional increase of sensitivity in small hawkmoth eyes. Our modelling provides an explanation for this counterintuitive finding: the relatively increased facet diameters decreased the amount of diffraction blur, thus benefiting spatial acuity in small eyes. Moreover, the observed scaling exponents optimised the eyes of large and small individuals to the smallest possible variation in sensitivity and spatial acuity, thus retaining a stable optical system across scales. Our results thus demonstrate that both visual functions are mutually optimised by scaling strategies in small superposition compound eyes.

## Results

To study how the eye size and eye morphology differed with body size in the superposition compound eye of *Macroglossum stellatarum* (Fig. 1A), we selected a total of 25 individuals with a wide range of body sizes (Fig. 1D). We obtained surface measures of their eyes (eye diameter: Fig. 1B,C, facet size: Fig. 2D) from light microscopy (9 animals) and X-ray microtomography (16 animals), which we combined in the subsequent analysis (see Methods). We relied on the X-ray tomography data for parameters requiring optical sections.

**Fig. 1.**
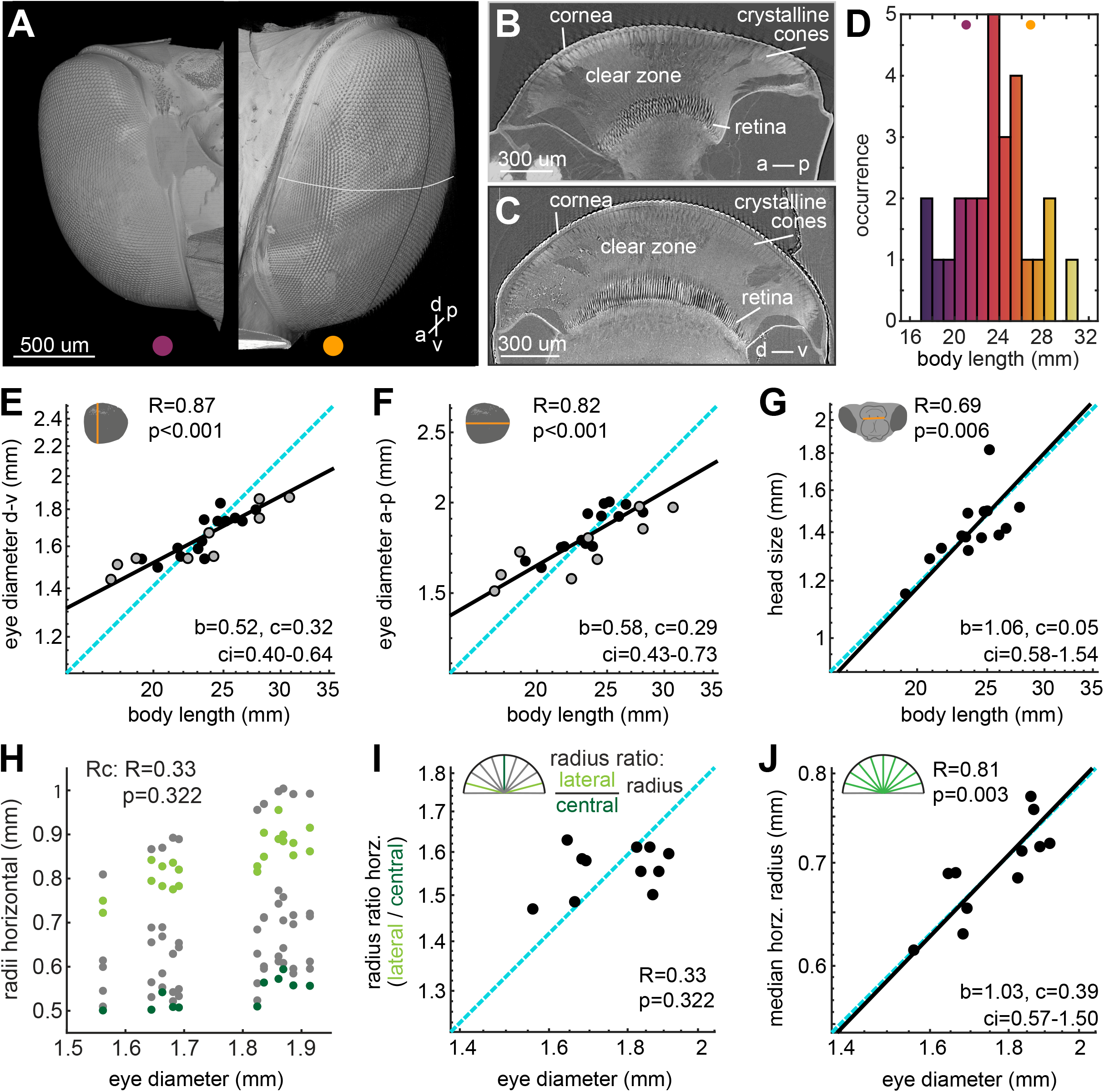
Allometric scaling of eye and head size in *Macroglossum stellatarum*. **A** X-ray microtomography images of two individuals of *M. stellatarum*. The animal to the left had a body length of 21.8 mm, the right one of 26.5 mm. The scale bar applies to both eyes. **B** Representative horizontal and **C** vertical section through the centre of the eye (see white and black lines in **A**) with the cornea, crystalline cones, clear zone and retina indicated. **D** Body length of the individuals selected for this study. Allometric scaling of the **E** dorsal-ventral, **F** the anterior-posterior diameter of the eye, and **G** the head size measured from the left to the right base of the mouth parts (Fig. S2A). **H** To test whether the shape of the eye differed across eye diameters, we measured the distance from the nodal point formed by the edges of the cornea to the corneal surface for nine evenly spaced radii in horizontal sections (see Fig. S3D-F for frontal ones). **I** We calculated the ratio between the average lateral radii (light green) and the central radius (dark green) as a proxy for the cornea’s shape, and assessed its allometric scaling. **J** The allometric scaling of the median of the central seven radius measurements (green) with eye diameter. **E**-**G**, **I**-**J** Data from individual hawkmoths was measured by either X-ray microtomography (black dots), or light-microscopy (grey dots). The dashed cyan line indicates isometric scaling and the black line represents the allometric scaling relationship. *R* is the Pearson correlation coefficient of the log-transformed data, and *p* denotes its statistical significance. Given the significant linear correlations in **E**-**G**, **J**, the allometric relationship was calculated using reduced major axis regression, with the exponential scaling exponent *b*, the normalization constant *c*, and the confidence interval *ci* of *b*.

**Fig. 2.**
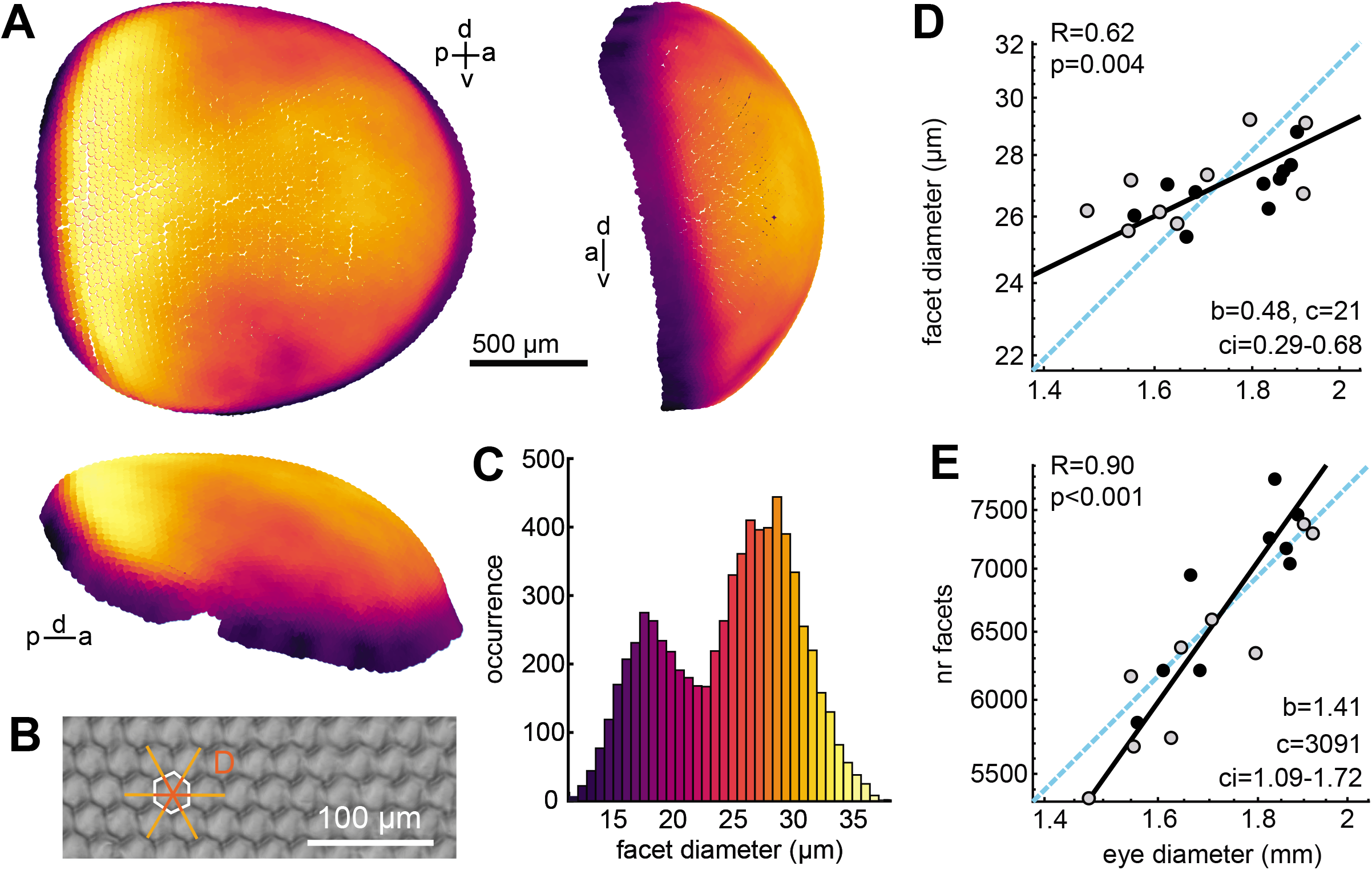
Cornea morphology and facet allometry of *Macroglossum stellatarum*. **A** 3D reconstruction of the facets of an example eye of *M. stellatarum* (total facets: 7111), with facet diameter indicated by the colour scale in **C**, top left: sagittal view, top right: anterior view, bottom: dorsal view. **B** The facet diameter was calculated as the average of 3 measurements (in light orange), to arrive at the facet distribution for the eye shown in **C**. **D** Allometric scaling of the median facet diameter of the eyes’ main facets (> 20 μm), and **E** the total number of facets with eye diameter. The total facet number was estimated by dividing the surface of the eye (approximated by a circular area based on the eye diameter) by the median facet size. **D**-**E** Data from individual hawkmoths was measured by either X-ray microtomography (black dots), or light-microscopy (grey dots). The dashed cyan line indicates isometric scaling and the black line represents the allometric scaling relationship. *R* is the Pearson correlation coefficient of the log-transformed data, and *p* denotes its statistical significance. Given the significant linear correlations in **D** and **E**, the allometric relationship was calculated using reduced major axis regression, with the exponential scaling exponent *b*, the normalization constant *c*, and the confidence interval *ci* of *b*.

### Eye size scales negatively allometric with body size

We observed significant negative allometry between eye diameter and body length with a scaling coefficient of 0.522 for the dorso-ventral eye diameter (Fig. 1C), and 0.577 for the anterior-posterior diameter (Fig. 1D). This indicated that smaller hawkmoths had relatively larger eyes than bigger moths. Moreover, the two axes of the eye had a highly significant correlation, which scaled isometrically (Fig. S3C), and allowed us to combine the eye diameter into a single measure where required, by taking the average of the two measures. Since the eyes comprise a substantial portion of the hawkmoth head, we also checked whether the scaling in eye size was mirrored by a scaling in head size. Since our specimen preparation did not preserve the entire head (see Methods), we measured proxies of head size using landmarks which could be reliably recognised in all preparations (Fig. S2A): the dorso-frontal (Fig. 1E) and lateral (Fig. S2B) extent of the mouth-part base, and the dorso-ventral extent of the head capsule surrounding the optic lobes of the brain (Fig. S2C). All of these scaled isometrically with body size, indicating that only the eyes of *M. stellatarum*, not the head as a whole, scale negatively allometrically with body size.

### Smaller animals have relatively larger, but fewer facets

Given the overall negative allometric relationship between eye and body size, we next investigated how structures of the eye that relate to spatial acuity and visual sensitivity scale with body and eye size. To quantify the size of the corneal facet lenses (Fig. 2A), we labelled all facets in two eyes, and 60-70 regularly spaced facets in all other eyes (n=19). The facet lenses varied in diameter across the hawkmoths’ eyes, with the largest facets being located in a median band along the anterior-posterior extent of the eye surface, and along the entire dorso-ventral extent of the posterior part of the eye. The histogram of all facet lenses of a completely reconstructed corneal surface clearly showed two peaks (Fig. 2B, Fig. S4A), representing the main facets of the eye and a ring of distinctly smaller facets located around the eyes’ perimeter, which are covered by scales in intact hawkmoths. The median diameters of outer facets, which might be structural in nature, did not correlate significantly with eye diameter (Fig. S4C). In contrast, the median diameters of the functional main facets (> 20 μm), correlated significantly with eye diameter (Fig. 2D) and body size (Fig. S4B).

A negative allometric scaling of facet diameter to eye diameter would indicate that smaller animals have fewer facets relative to their eye diameter than large ones – provided that the relationship between the surface area of the cornea and eye size did not differ. Since the surface area depends on the shape of the cornea, we analysed the scaling of the cornea’s curvature with eye diameter (Fig. 1H-J, Fig. S3D-F). We calculated the ratio of the central and lateral radii of the eye in horizontal (Fig. 1I) and frontal sections (Fig. S3E) at the dorso-ventral and anterior-posterior median of the eye, respectively. There was no significant correlation of the curvature ratio with eye diameter (Fig. 1I, Fig. S3E), while the average radius of the cornea scaled isometrically with eye size (Fig. 1J, Fig. S3F), indicating that the corneas’ curvature remained the same in large and small eyes. This confirmed the validity of our approach to estimate eye surface based on eye diameter. It also allowed us to estimate the total number of facets per eye, by dividing the eye surface area by the median facet diameter. The total facet number scaled positively allometric (Fig. 2E), with the lower-bound confidence interval exceeding isometry. Thus, smaller hummingbird hawkmoths invested in larger facet diameters at the cost of the total number of facets.

To assess whether the shape of the cornea differed with eye diameter, we measured evenly spaced eye radii in horizontal and vertical sections (Fig. 1H, S3D). The ratio of the two lateral-most radii and the central one in each section was used as an indicator for shape: if, for example, a larger eye was rounder than a smaller one, the ratio would be smaller in larger eyes, while it would remain the same, if the shape of the cornea did not change. We thus analysed the allometric scaling of the radius ratio with eye size, and found there was no significant correlation in either the horizontal (Fig. 1I) or frontal sections (Fig. S3E). The median of all radii in both horizontal and frontal sections scaled isometrically with eye diameter (Fig. 1J, S3F)

### Rhabdom distance, but not length, scales negatively isometric with eye size

We next analysed whether the scaling relationship of facet lenses transferred to the retina. In a typical apposition compound eye, each facet lens forms a structural unit with a group of photoreceptors (the rhabdom), termed an ommatidium. In most superposition compound eyes, the 1:1 relationship between facet lenses and photoreceptive units exists as well, although the optical relationship is uncoupled by the optical units in the superposition pupil focusing light from many facet lenses onto a single rhabdom (Exner, 1891; Warrant & McIntyre, 1993). In the hummingbird hawkmoth, the anatomical 1:1 relationship between facet lenses and retinal units was called into question, due to an optically measured inhomogeneity in facet diameter and retinal packing (Warrant 1999). Since the tracheal sheaths surrounding the photoreceptors (Warrant et al., 1999) provided high optical contrast, we could fully reconstruct all rhabdom positions in two eyes (Fig. 3C, inset). From this, we calculated inter-rhabdom distances (IDR, Fig. 3C, inset) similar to the inter-facet distances (Fig. 2A). The inter-facet and inter-rhabdom distances showed very different local patterns across the eye, highlighting that the facet distribution was uncoupled from the retinal one. However, the total number of rhabdoms and facets identified in two eyes were very similar. Indeed, the number of rhabdoms was 6% and 8% higher – a divergence likely caused by an underestimation of the number of facets, as some of the structural facets could not be resolved.

**Fig. 3.**
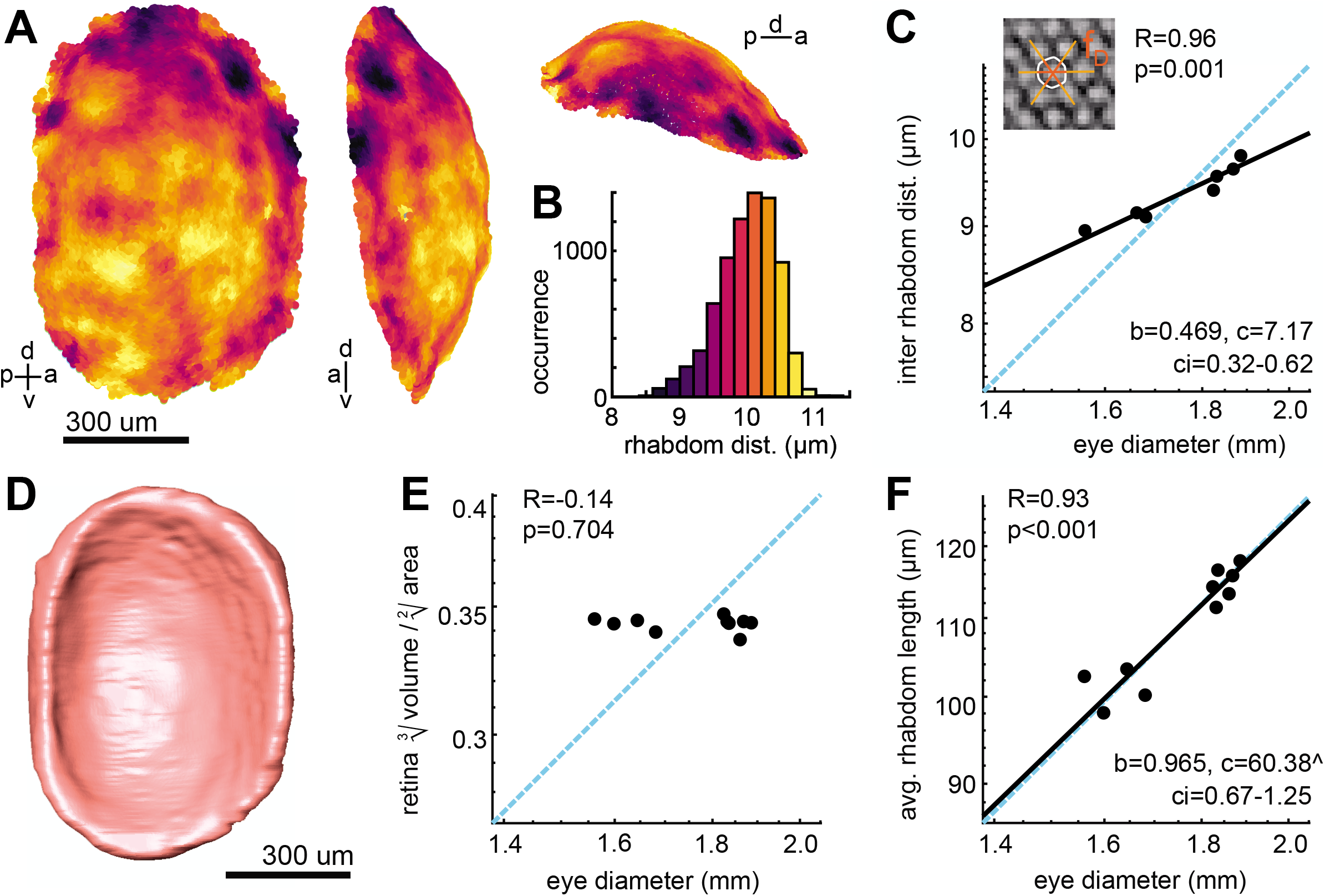
Retinal morphology and rhabdom scaling. **A** 3D reconstruction of the rhabdoms in an example retina (with 7560 rhabdoms), with inter rhabdom distance (IRD) indicated by the colour scale in **B**, top left: sagittal view, top right: anterior view, bottom: dorsal view. **C** The inter rhabdom distance (IRD) was measured as the average distance between rhabdoms as shown in the inset. Allometric scaling of the IRD. **D** 3D-reconstruction of the retinal volume of the example eye. **E** Differences in retinal shape assessed as the ratio between retinal volume and surface area across eye diameters. **F** A conserved retinal shape across eye sizes allowed us to use the thickness of the retina as a proxy for the average rhabdom length, calculated as the volume divided by half the surface area. **C**,**E**,**F** Data from individual hawkmoths was measured by X-ray microtomography (black dots). The dashed cyan line indicates isometric scaling and the black line represents the allometric scaling relationship. *R* is the Pearson correlation coefficient of the log-transformed data, and *p* denotes its statistical significance. Given the significant linear correlations in **C**,**F**, the allometric relationship was calculated using reduced major axis regression, with the exponential scaling exponent *b*, the normalization constant *c*, and the confidence interval *ci* of *b*.

For all eyes, we determined the average IRDs in the centre of the retina (see Methods) as a measure for the separation of the anatomical sampling base of the eyes. This IRD showed only a single-peaked distribution (Fig. 3B), as compared to the double-peaked distribution of the facet sizes. Nevertheless, there was still considerable variation in the IRDs (Fig. 3B), which was systematically larger in the ventral than the dorsal half of the retina (Fig. 3A). The rhabdom distance scaled negatively allometric with eye size across individuals (Fig. 3C), indicating that smaller individuals had distinctly larger IRDs than expected for their eye size. Moreover, IRDs scaled with the same coefficient as facet diameter across eye size (Fig. 2D), and indeed there was a linear relationship between IRDs and facet diameters (Fig. S6C), giving additional support to the notion that that the number of photoreceptor units in the retina matches the number of facets in the cornea.

To assess how rhabdom length scaled with eye size, we used the thickness of the retina as a proxy. This is possible if the retinal shape was the same in animals of different body size. We confirmed this by the comparing the ratio of retinal volume and surface area across eye sizes (Fig. 3E): if the retina became flatter with eye size, the ratio should decrease, while it should increase if the retina became thicker. Since the ratio remained the same across eye size (Fig. 3D), we concluded that retinal shape did not scale with eye size. We thus estimated the rhabdom length by dividing the retinal volume by half its surface area. Unlike IRD, rhabdom length scaled isometrically with eye size (Fig. 3C). Thus, smaller hummingbird hawkmoths invested in larger IRDs at the cost of total number of rhabdoms, while the length of their rhabdoms scaled isometrically with size.

### Both sensitivity and spatial acuity are optimised in small hawkmoths

To understand how the scaling of the optical and sensory structures affect the function of large and small hawkmoth eyes, we used the observed allometric relations to calculate key performance measures of eyes: single-ommatidium sensitivity (Fig. 4A, Methods: equation 5, according to (Warrant, 1999)), spatial resolution as the photoreceptor acceptance angle (Fig. 4B, Methods: equation 4, according to (Land et al., 1997)), and the limiting feature of spatial acuity: the half-width of the Airy disc (Fig. 4C, Methods: equation 3). To do so, we used the measured scaling coefficients of the facet diameter, inter-rhabdom distance (IRD), and rhabdom length to estimate eye performance for animals of different body lengths. We approximated the scaling of rhabdom diameters by the scaling of the IRD, assuming that the tracheal sheath surrounding each rhabdom (which contributed to the IRD, but is not optically functional), scales isometrically with eye size and remains constant across the eye, which electron microscopic sections support (Warrant et al., 1999).

**Fig. 4.**
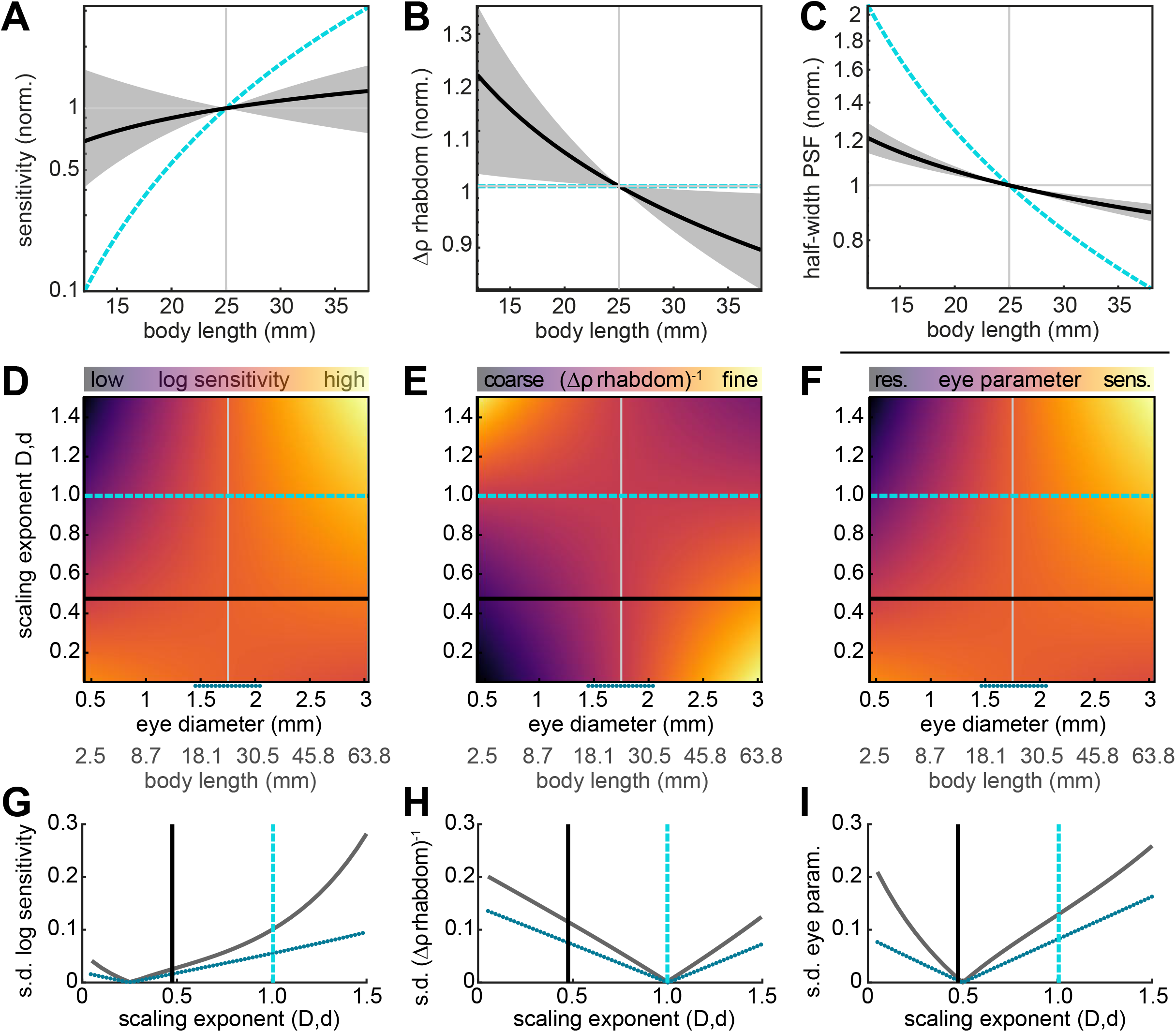
Model estimation of the allometry of spatial acuity and sensitivity. We used the measured allometric relations of the inter-facet and inter-rhabdom distance *D* and *d*, rhabdom length *l* and focal length *f* (the latter two scaling isometrically) to calculate **A** the sensitivity of a single ommatidium according to Warrant & Nilsson, 1998), **B** the rhabdom acceptance angle and **C** half-width of the point spread function (PSF), according to Land et al., 1997). All estimates of eye performance were calculated for body lengths ranging from 12.5 to 37.5 mm, and normalised to a median sized animal of 25 mm body length. All calculations were compared to estimates based on an eye in which all parameters scaled isometrically (cyan line). The confidence intervals were computed by applying the same calculations to the scaling parameters (exponent and Y-axis intercept) with added and subtracted confidence intervals obtained from the regression analysis. **D** Log-transformed sensitivity, **E** rhabdom acceptance angle, and **F** eye parameter were calculated for a range of scaling exponents applied to the inter-facet and inter-rhabdom distance *D* and *d* (see Methods). **D**-**F** The resulting values are depicted for different eye diameters (with corresponding body lengths indicated in grey), normalised to the largest sensitivity, smallest rhabdom acceptance angle, and largest eye parameter. The measured scaling exponent is indicated by the black line, and isometry by the blue dashed line. The dotted blue line below the x-axis indicates the measured size variation. Variation in **G** log sensitivity, **H** rhabdom acceptance angle, and **I** eye parameter for a given scaling exponent across eye sizes, quantified as the standard deviation (s.d.) for the entire range of eye diameters (grey line), and the measured range (blue dotted line). The black line indicates the measured exponents, and the blue dashed line isometry.

For these calculations, the focal length of the eye was also required. Although it cannot be directly determined anatomically in aspherical superposition compound eyes (Warrant, 1999), we could show that it is valid to apply the same scaling coefficient for the focal length as for the eye diameter. The focal length in superposition compound eyes can be measured as the distance from the eyes’ nodal point to the tip of the retina (Land et al., 1997; Snyder, 1977). The nodal point is determined by the eyes’ radius, which scaled isometrically with eye size (Fig. 1J). The distance from the nodal point to the tip of the retina is determined by the eye radius, and the distance of the retina to the cornea. The latter remained constant across animals of different sizes (Fig. S5), suggesting that the scaling relationship of the focal length is determined by the scaling of eye size.

Given these scaling parameters, we could show that the sensitivity of a single ommatidium scaled with a distinct negative allometry compared to an eye in which all structures scaled isometrically (Fig. 4A): with isometric scaling, each ommatidium of an animal with 12.5 mm body length would have a ten times reduced sensitivity compared to an animal with 25 mm body length (the median). The reduction in sensitivity given the measured scaling was only 30%, and thus seven times higher than for isometric scaling. Moreover, the 95% confidence intervals still included the sensitivity value of the median sized animal, indicating that there is a neglible difference in sensitivity between animals differing in size by a factor of 2.

For the estimate of spatial resolution, our model showed that the photoreceptor acceptance angle of animals of 12.5 mm body was 20% larger compared to the median animal of 25 mm body length – with confidence intervals not overlapping the median (Fig. 4B). This represented a distinct difference from isometric scaling, which did not predict any differences from a median sized animal, because both the rhabdom diameter and focal length scaled isometrically in this case. The optical limitation of spatial resolution, the half-width of the fundamental mode of the point spread function (PSF) which causes diffraction at a single facet lens (see equation 4, Methods), scaled so that smaller eyes had a relatively smaller diffraction blur circle than they would have had with isometric scaling (Airy disc, Fig. 4C).

### The scaling of facets and rhabdoms minimised differences in the eyes’ optical function across body sizes

We next assessed how the observed scaling exponents of the inter-facet and rhabdom distance determined the performance of eyes across sizes, compared a range of hypothetical scaling exponents representing negative and positive allometry, as well as isometry. We focused on these two structures, because they diverged strongly from isometry with eye size and scaled with very similar exponents, so that a common exponent could be assumed for modelling (0.48 for facet diameter, 0.47 for rhabdom diameter, average of 0.475 indicated as the black line in Fig. 4D-F). The focal and rhabdom lengths, which also contribute to the acuity and sensitivity of the eye, scaled isometrically with eye size. We calculated the ommatidial sensitivity and rhabdom acceptance angle as before, across a range of possible allometric scaling parameters for a range of eye sizes (Fig. 4D-E). We also performed this calculation for the eye parameter (Fig. 4F, Methods: equation 6), a measure of the eyes’ optimisation for sensitivity or spatial acuity (smaller values suggest optimisation for acuity, large values for sensitivity).

For ommatidial sensitivity, isometric or positive allometric scaling resulted in distinctly higher sensitivity in larger than in smaller eyes (Fig. 4D). This strong divergence decreased down to a scaling exponent of approximately 0.3, below which the sensitivity was moderately higher in smaller than larger eyes. The observed scaling exponents of 0.48 for the facet diameter and 0.47 for the rhabdom diameter (average of 0.475 indicated as the black dashed line in Fig. 4D-F) yielded a moderate difference in sensitivity across eye sizes, as also described in Fig. 4A. A very different performance for small and large eyes was obtained for the rhabdom acceptance angle, where larger animals would have coarser angles than smaller ones for a scaling exponent above 1, and vice versa below 1. The same acceptance angle was predicted for all eye sizes with isometric scaling (Fig. 4E). Finally, the eye parameter, flipped in its effect for smaller and larger eyes at scaling exponents close to those measured in hawkmoth eyes (Fig. 4F): for scaling exponents higher than 0.5, larger eyes are optimised more strongly the sensitivity, and this was also the case for smaller eyes for scaling exponents below 0.5. Across all three eye performance values, the scaling exponents observed in the eyes of *M. stellatarum* reduced the variance in sensitivity and eye parameter across eyes of different sizes compared to isometric scaling (Fig. 4G, I): the observed scaling exponents were close to the overall minimum of variance across eyes for sensitivity (Fig. 4H), while they fell right into the minimum for the eye parameter (Fig. 4I). This indicates that the scaling of facet and rhabdom diameters in the superposition compound eyes of hummingbird hawkmoths are optimised to reduce the variance in eye performance across eye and body sizes.

## Discussion

In this study, we used 3D X-ray microtomography to provide the first quantification of allometric scaling of the morphological and functional features of a superposition compound eye. We revealed that the overall scaling of the hummingbird hawkmoth’s eye with body size was negatively allometric, as in many other insects. Even though the superposition optics provides a generally higher sensitivity to light than the optics of apposition compound eyes of a similar size, we found that non-isometric scaling reduced the loss in sensitivity in the smaller eyes of smaller individuals even further. Overall, the allometric scaling of the hawkmoths’ eye parameter minimises differences in absolute sensitivity and spatial acuity across eye and body sizes.

### Local inhomogeneities in hummingbird hawkmoth superposition eyes

To quantify the allometric scaling of hummingbird hawkmoth superposition eyes, we undertook the first 3D structural characterisation of these eyes, which revealed some unexpected features of their visual system. It has been described previously that, unusually for optical superposition compound eyes (Exner, 1891; Meyer-Rochow & Gál, 2004), hummingbird hawkmoth compound eyes are inhomogeneous (Warrant et al., 1999). Unlike the spherical eyes of their nocturnal relatives (for example *Deilephila elpenor*, (Stöckl et al., 2016b)), their cornea and retina are locally flattened, particularly in the anterior-posterior axis. Furthermore, facet and rhabdom diameters are inhomogeneously distributed across the eye (Figs. 2,3), reminiscent of the local acute zones in apposition compound eyes (Land & Eckert, 1985; Land, 1989; Straw et al., 2006; Taylor et al., 2019)). Our results confirmed previous data obtained using tissue sections of a band of increased facet diameter along the lateral midline of the eye (Warrant et al., 1999). In addition, we revealed that the largest facets in the hawkmoth eye are positioned at the posterior edge of the eye, extending over the entire dorso-ventral axis. These facets were nearly 30% larger than the average facet diameter across the eye, suggesting that increased sensitivity in the posterior visual field is of high importance to the hawkmoths. This might serve to recognise approaching predators as early as possible, especially while hawkmoths are at their most vulnerable, hover-feeding from flowers (Stöckl & Kelber, 2019; Wasserthal, 1993). Our data also provides evidence for two classes of facets in the eye of hummingbird hawkmoths: the main facets of the eye, and a group of distinctly smaller facets around its perimeter (Fig. 2A) that are covered in scales in intact hawkmoths. These two groups are visible as two clear peaks in the facet diameter histograms (Fig. 2C, S.4A). The fact that the small perimeter facets did not scale with eye size (Fig. S4C), while main facets did (Fig. 2D), further suggests they are unlikely to be optically functional, but instead have a structural role. More research into the optical axes and focussing properties of these small facets will be required to elucidate whether they do play a functional, or a purely structural role.

### Anatomical existence of ommatidia in hummingbird hawkmoth eyes

Unexpectedly, our findings call into question an interpretation of previous anatomical findings from hummingbird hawkmoth eyes, namely the suggestion they lack true ommatidia in the developmental and functional sense, because rhabdom density is up to four times higher than facet density in the frontal acute zone (Warrant et al., 1999). In the retinas in which we fully reconstructed the positions of all rhabdoms, we did not observe this effect (Fig. 3). On the contrary, rhabdoms were spaced more widely in the fronto-ventral part of the eye than the dorsal hemisphere (Fig. 3A). The close match of identified facets and rhabdoms in the fully reconstructed eyes suggests that anatomically, although not necessarily functionally, the optical and receptive elements form a single unit in the eye of hummingbird hawkmoths. The denser rhabdom packing in the frontal eye observed previously using opthalmoscopic measurements might thus have been an optical effect. The rounded frontal cornea focusing light onto a very flat frontal retina could potentially produce a magnification of the focused image, leading to increased spatial resolution without a denser rhabdom packing. Future optical modelling will have to reveal whether this hypothesis holds, while developmental investigations might unravel how the highly inhomogeneous distribution of facet and rhabdom mosaics emerges.

### Scaling of eye size compared to other insects

The scaling of the superposition eyes of hummingbird hawkmoths followed the same general trend described for the apposition eyes of other insect groups: they scaled negatively allometric with body size (bees: (Jander & Jander, 2002; Spaethe, 2003; Streinzer & Spaethe, 2014; Taylor et al., 2019), ants: (Perl & Niven, 2016a; Zollikofer et al., 1995), butterflies: (Merry et al., 2006; Rutowski, 2000), and flies (Currea et al., 2018). The scaling exponent we observed in hawkmoths (average: 0.55) was slightly larger than in bumblebees (0.45, (Taylor et al., 2019)), and fell well within the ranges described for ants (Perl & Niven, 2016a) and fruit flies (Currea et al., 2018). Interestingly, head size scaled isometrically in the hawkmoths, thus resulting in proportionally smaller heads than eyes in smaller individuals. In line with this, overall brain size and optic lobe size also scales isometrically in this hawkmoth species (Stöckl et al., 2016a), suggesting separate growth regulation for head and brain size on one hand, and eye size on the other hand.

The comparison of morphological structures related to visual sensitivity between hawkmoths and previously studied insects is of particular interest, since the hawkmoths’ superposition compound eyes provide high visual sensitivity due to its specialised light-collecting optics (Exner, 1891; Warrant & Nilsson, 1998). We hypothesised that the trend to larger sensitivity in larger apposition compound eyes, as seen in bumblebees (Spaethe, 2003; Taylor et al., 2019), would be less pronounced in the hummingbird hawkmoth, where sensitivity might be under less selection pressure because the superposition pupil increases light capture by 200-times (Stöckl et al., 2017c; Warrant et al., 1999). Surprisingly, the opposite was the case: the allometric scaling exponent of the facet diameter with eye size was distinctly smaller than in bumblebees (0.71 (Taylor et al., 2019)) and smaller than in fruit flies (0.57 (Currea et al., 2018)). The consequence of the relatively increased facet and rhabdom diameters, in combination with isometrically scaling focal and rhabdom lengths, was a distinctly increased ommatidial sensitivity in smaller eyes compared to isometric scaling (Fig. 4A). Thus, compared to insects with less light-sensitive apposition eyes (Currea et al., 2018; Spaethe, 2003; Taylor et al., 2019), the highly sensitive superposition compound eyes of hawkmoths had the strongest optimisation for single-ommatidia sensitivity.

### Benefits of relatively increased facets and rhabdoms in superposition eyes

While the investment in high sensitivity might seem counterintuitive, one needs to consider that increased facet and rhabdom diameters do not just support ommatidial sensitivity, but can also improve spatial acuity if the eye is diffraction limited (Land et al., 1997; Snyder, 1977; Snyder et al., 1977). The strongly negative allometric scaling of the facet diameter would reduce the size of single-facet based diffraction blur compared to isometric scaling (Fig. 4C). This scaling also results in relatively increased rhabdom diameters in small individuals, which further limits potential light-leakage effects into neighbouring ommatidia due to wave-guiding in the rhabdoms (Warrant et al., 2007), because the rhabdom diameters remain several times larger than the wavelength of visible light (Fig. 3C). Light leakage is further prevented by the tracheal sheet around each photoreceptor unit (Warrant et al., 1999).

While previous work suggests that the diffraction blur caused by a single facet in a compound eye linearly adds to the photoreceptor acceptance angle (Snyder, 1979), and thus compromises spatial resolution, this assumption does not seem to hold for superposition compound eyes (Stavenga et al., 2006), nor indeed for apposition compound eyes (Stavenga, 2003; Warrant & McIntyre, 1993). In superposition compound eyes, the interaction of partially coherent light waves focused on a single rhabdom causes complex diffraction patterns that depend on the number of ommatidia in the superposition pupil (Stavenga et al., 2006). This effect decreases the extent of the blur resulting from diffraction, and might thus release superposition compound eyes from the diffraction limitations on spatial acuity that are imposed by single facets. If this was indeed the case for hummingbird hawkmoth eyes, which future optical modelling studies need to confirm, the relatively enlarged facets in smaller hawkmoths might not contribute to improved spatial acuity by decreasing the half-width of the diffraction blue compared to isometric scaling (Fig. 4C).

It is furthermore important to consider that visual sensitivity does not just set the absolute detection limits of the eye, but also determines how fine contrasts a visual system can resolve (Land et al., 1997; Snyder, 1977). Thus, while sensitivity is high due to the eye design in hawkmoths, and these diurnal insects can still see (Stöckl et al., 2017c) and perform visual behaviours even at moonlight intensities (Stöckl et al., 2017b), the observed scaling might serve to maximise sensitivity for the purpose of retaining high contrast resolution in small hawkmoths. One benefit of high contrast sensitivity even for diurnal insects is the detection of small objects, which is ultimately restricted by the sensitivity of individual photoreceptive units (Rigosi et al., 2017). Furthermore, high contrast sensitivity paired with high spatial resolution might be particularly adaptive for hovering insects, as it allows them to resolve motion cues both at slow hovering and fast forward flight speeds (O’Carroll et al., 1996). Thus, allometric scaling of facets and rhabdoms to retain high contrast sensitivity in small hawkmoths might provide benefits for spatial and motion tasks, on top of the high absolute sensitivity that their superposition compound eyes provide.

### Optimising eye performance across scales

One striking hypothesis for the scaling of the different optical structures emerged when we assessed how the observed scaling affected the performance of the hawkmoth eye compared to other possible scaling coefficients. The measured scaling exponents reduced the variation in sensitivity and spatial acuity across eye sizes, compared to isometric scaling. Indeed, they optimised the eye parameter very close to the minimum in variation across scaling factors, meaning that the eyes of larger and smaller hawkmoths varied the least possible in their spatial acuity and sensitivity (Fig. 4). This likely benefits the subsequent processing of information from the eyes, because processing strategies can be largely retained across size ranges – particularly with respect to the processing that affects spatial resolution and visual sensitivity (Stöckl et al., 2017a; Stöckl et al., 2020; Warrant, 1999). As discussed above, scaling that changes the contrast and spatial properties of the visual system might alter the perceptual thresholds for object or motion detection, for example, and thus require subsequent adjustments in the visual circuits to enable individuals of different sizes to successfully perform visual behaviours. Motion vision provides an interesting case, because the spatial and temporal properties of the visual input are tightly entwined in the motion percept (Borst & Egelhaaf, 1989). Consider, for example, two hummingbird hawkmoths with different body sizes, and thus with different spatial acuity due to allometric scaling, flying at the same speed in the same environment. Their neuronal responses to motion will be different, because motion-sensitive neurons are temporal frequency tuned, and the temporal frequencies they observe will differ depending on the spatial sampling base of the eye (Borst & Egelhaaf, 1989). How then, would the motion vision system be adjusted to optimally code motion in the velocity range these insects experience – or does the adjustment take place on the behavioural side, so that moths with higher spatial acuity fly at lower speeds than those with lower acuity? Scaling the eye so that changes in spatial acuity and contrast sensitivity are minimised between large and small individuals, as observed in the hummingbird hawkmoths, will minimise the need for such behavioural or physiological adjustments, and thus markedly simplify the subsequent visual processing across body size ranges.

### Adaptive consequences of eye scaling in solitary and social insects

The reduction of variation in sensitivity and acuity across hawkmoth sizes also suggests that larger and smaller hawkmoths would have similar visually-driven behavioural abilities. In terms of spatial acuity, this is supported by recent findings, which show no difference in spatial resolution between large and small hawkmoths in an optic flow task (Grittner et al., 2021). Given that the estimated decrease in the photoreceptor acceptance angle in the smallest tested hawkmoths was 15% lower than that in an 80% larger moth (Fig. 4B), and the range of spatial frequencies the hawkmoths responded to behaviourally (Grittner et al., 2021), the lack of a behavioural phenotype might not be surprising. This is in stark contrast to bumblebees, where the spatial resolution improved by 30-50% (measured as the inter-ommatidial angle) in 50% larger bumblebees. This distinct scaling of visual sensitivity with body size manifests in behaviour: larger bees forage at lower light intensities (Kapustjanskij et al., 2007) and detect smaller point-targets than smaller ones (Spaethe, 2003; Streinzer et al., 2016). In general, there might be a higher tolerance for variations in eye performance across scales in social insects, since the unit of selection is the colony (Korb & Heinze, 2004), not the individual. In bumblebees, the workers that leave the nest to forage are typically larger individuals (Cumber 1949), so that a scaling of sensitivity benefits the colony in foragers with a higher sensitivity, while the smaller individuals can take up other tasks in the colony. In hawkmoths, where the unit of selection is the individual, a strong scaling of visual sensitivity with eye size would be mal-adaptive to a distinct proportion of the population, which might therefore have a lower tolerance for performance scaling with eye size. Future comparative work is required to resolve which role solitary lifestyle, phylogenetic heritage and eye design play in the allometric scaling we found in hummingbird hawkmoths.

### Conclusion

Insect compound eyes provide an ideal model to study how miniature optical systems optimise their performance across scales. In this study, we provide the first quantification of the allometric scaling of the morphology and functional characteristics of a superposition compound eye. We revealed that this eye type follows the same trend for negative allometry of eye size with body size as many other insects. Our results demonstrate how eye scaling benefits the performance of the eye in terms of sensitivity and spatial resolution. By showing that the measured scaling factors in hummingbird hawkmoths minimise the variation in eye performance across eye sizes, we open the field for future investigations into how allometric scaling optimises eye performance in different species and optical systems.

## Methods

### Animal measurements

Hawkmoths (males and females) were kept on a 14:10h light/dark cycle in flight cages (60 cm x 60 cm x 60 cm) and fed with artificial feeders (Pfaff & Kelber, 2003) that contained a 20% sucrose-water-solution for several days before being used in experiments. To investigate the scaling of eye morphology, we selected a total of 25 individuals with a wide range of body sizes (Fig. 1D). We weighed all animals before the preparation of eyes, and photographed every animal to determine their body and wing size using the Fiji software (Schindelin et al. 2012). Total body length was measured from their anterior to posterior extent, the thorax width was measured from wing-base to wing-base, the total wing length was measured from the base to the tip of the wing for both wings and averaged, and the inner wing length was measured from the base to the inner turning point of the wing (see Fig. S1 for descriptions of all measurements). For most comparisons, we relied on the body length as a measure of body size, as this had the highest correlation with other body size parameters, such as the weight, and the size of the wings (Fig. S1). Some data in this study did not include body length measurements, but only outer wing length. To obtain an estimate of body length we used the highly linearly correlated relation between body and wing length in hummingbird hawkmoths (Fig. S1, see also (Kihlström et al., 2021)), by computing the allometric scaling between the two parameters (see *Allometry calculations*) and solving the equation for body length.

### Head and eye preparations

We prepared the eyes of 16 hawkmoths for microtomographic imaging. For that, we retrieved the heads of cold-anesthetized hawkmoths, and removed their antennae and the dorsal part of their head capsule, as well as the mouth parts, with a sharp razor blade. We furthermore removed the lateral tip of the left eye to improve the impregnation of the sample with fixative solution and resin. This was particularly important since the large clear zone of the superposition compound eyes, and the high amount of trachea in and around the brain and retina posed a considerable challenge for fixation and embedding of the tissue. We immediately fixed the dissected samples in 3% paraformaldehyde, 2% glutaraldehyde, and 2% glucose in phosphate buffer (pH ~7.3, 0.2M) for 3 hr, and then washed these in phosphate buffer before immersing them in 2% OsO4 for 1 hr to enhance the X-ray absorption contrast (Ribi et al., 2008). After washing in buffer again, the samples were dehydrated with a graded alcohol series, and acetone was used to transition the samples to epoxy resin (Agar 100, Agar Scientific) in multiple steps of increasing concentration (see (Stöckl et al., 2016b)). The samples in wet epoxy were placed with their dorsal side facing up on cured Epoxy mounts and cured in an oven at 60°C for ~48 hr.

### X-ray microtomography imaging

Tomographic imaging of moth heads was performed at the Diamond-Manchester Imaging Beamline I13-2 (Pešic et al., 2013; Rau et al., 2011) at the Diamond Light Source UK (proposal MT13848). Fixated heads were imaged using 4 x total magnification (with an effective pixel size of 1.625 μm) using a pco.edge 5.5 (PCO AG) detector with a 50 mm propagation distance. For further details, see Taylor et al. (2019).

### Eye measurements

The scan data was compressed from 32 bit to 8 bit, and cut to contain only the region of interest, using Dristhi (Limaye, 2012). The subsequent data analysis was performed on reconstructed 3D volumes in Amira (Release 6.8, Thermo Fisher Scientific). Data was extracted only from specimen that had sufficiently high quality of preparation and scanning to reliably extract the following measures. The *Source Data* file provides an overview of which data was extracted for each specimen. Using the *3D measurement* tool, we extracted the anterior-posterior and dorso-ventral diameters of each eye (Fig. 1E,F, orange lines), as well as three measures of head size: the dorso-ventral and lateral extents of the mouth-part base, the dorso-ventral extent of the right optic lobe (Fig. S2A). A subset of eye diameters (9 animals, highlighted in the respective figures) was measured by photographing the eye laterally through a stereoscope with a scale bar.

The median facet diameters in this subset of data were determined from corneal imprints with nail polish (Stöckl et al., 2016b), while all other facet diameters were measured on the 3D volumetric data in Amira: to this aim, the data was rendered using the *isosurface* tool with an individually adjusted brightness histogram, to optimally resolve the surface of the eye (for example Fig. 1A,B). Then, 60-70 measurements spaced in regular distances over the entire eye were performed. Each of these measurements comprised a group of seven facets (one central facet and its six neighbours), for which three measurements were performed, spanning from the outer edge of a facet to the opposing facet beyond the central one (see Fig. 2B, and (Taylor et al., 2019)). The results were averaged and divided by the number of facets spanned (three), to obtain an average measure of the facet diameter at each of the 60-70 positions on the retina. These measurements formed the basis for the inter-facet-distance histograms (Fig. 2C, S4A). In addition, we also reconstructed the positions of all corneal facets in two selected eyes and used their coordinates to calculate the inter-facet distances for all facets in these eyes (Fig. 2A). Both the complete reconstruction, as well as the lower resolution sampling of facet distances revealed a bimodal distribution of facet diameters (Fig. 2C, Fig. S4A). The two peaks of the distribution represent the main facets of the eye and a ring of distinctly smaller structural facets around the eyes’ perimeter, which are covered by scales in intact hawkmoths. In the subsequent allometric analysis we separated the two facet groups using a cut-off of 20 μm, because their scaling with eye size differed.

We used the same measurement strategy to measure the inter-rhabdom distances in the retina. To this aim, we virtually removed the distal portion of the eye to reveal the distal surface of the retina in the lateral eye (where rhabdoms were most clearly separable). Here, we performed 20 measurements (containing three measurements each as for the facet distances), which were spaced 7-8 rhabdoms apart. We calculated the inter-rhabdom distances as for inter-facet distances (Fig. 3C). In the two eyes in which all facet positions were reconstructed, we virtually exposed the entire distal surface of the retina and reconstructed the position of every resolvable rhabdom (Fig. 3A,B).

We further analysed a variety of functional parameters on optical sections through the anterior-posterior median (horizontal section, Fig. 1B) and dorso-ventral median (frontal section, Fig. 1C) of the eye. For each eye and section orientation, we conducted 10 evenly spaced measurements of cornea thickness (Fig. S3B), crystalline cone length (Fig. S3C), and distance between retina and cornea (Fig. S5A,D). To assess whether the curvature of the eye differed with eye size, we performed nine evenly spaced measurements of the eye radius, whose medial border was defined by the medial edge of the cornea. The nodal point for the measurements was placed in the centre of this line, forming the first two measurements. From there, a measurement perpendicular to the medial measurements was performed to the distal-most extent of the cornea, and three further measurements were spaced evenly between these on both sides (Fig. 1I, S3E). Using these, we calculated the ratio between the middle radius, and the two lateral-most ones next to the edge radii (Fig. 1H, S3F). If the overall curvature of the cornea differed, for example if the eye became flatter as animals became larger, the ratio should decrease, as the central radius would decrease in length relative to the lateral ones. To assess potential difference of cornea curvature with eye size, we computed the allometric scaling of the curvature ratio with eye diameter.

Finally, we calculated an estimate of the number of facets per eye as the corneal surface area divided by the facet diameter. Since we were not able to reconstruct the surface area of all eyes, we derived a scale factor from the eight eyes with fully reconstructed corneal surfaces that allowed us to estimate the eye surface area from the eye diameter – which was possible since the overall curvature of the cornea did not differ (Fig. 1H, S3F):

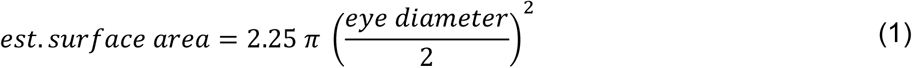

We calculated the number of facets per eye as the estimated surface area divided by the median functional facet size. This measure does not take into account the number of structural facets but provides a conservative estimate for the scaling of the functional facets in the eye.

### Allometry calculations

To assess the scaling relationship between different eye and body sizes, we first tested for a significant linear correlation between the two log-transformed parameters in questions, by means of the Pearson correlation coefficient (*R*). Only if a significant log-linear correlation was found (p<0.05), we proceeded to test the allometric scaling. For this, we used Model II (reduced major axis) regression implemented in the *gmregress* script for MATLAB (A. Trujillo-Ortiz, www.mathworks.com/matlabcentral/fileexchange/27918-gmregress, retrieved March 19, 2020). This provided the scaling exponent *b* and the scaling constant *a* of the allometric relationship

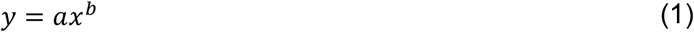

Fitted to the parameters in the log-transformed version of the equation:

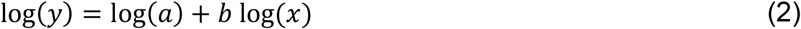

The scaling exponent *b* describes the slope of the linear relationship, and the log-transformed scaling constant *a* describes the y-axis intercept (Warton et al., 2006).

### Acuity and sensitivity calculations

To understand how the scaling of the optical and sensory components of hawkmoth eyes contribute to their function, we used the measured allometric relations of eye structures to calculate how the diffraction limit, acceptance angle and sensitivity of a single ommatidium scale with body and eye size of the hawkmoths. To this aim, we calculated how these measures differed relative to an average individual with 25 mm body length (Fig. 4). We applied all parameters in the following formula, which are not constant for animals of varying body sizes, and constants where necessary to retain proportionality, and scaled their values according to the measured allometric relationships with body size. To obtain confidence intervals for these calculations, we applied the upper and lower confidence intervals of each scaling exponent and intercept to obtain an estimate of the lowest and highest value of each eye parameter for each animal size.

Since the rhabdom diameter estimated from the inter-rhabdom distance and previously measured by (Warrant et al., 1999) distinctly exceeded the average wavelength of visible light, we calculated the half-width of the point-spread function *hw_PSF_* (Airy disc) created by the optics of each facet (Fig. 4C) as

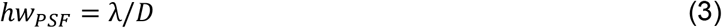

according to (Land et al., 1997; Warrant et al., 2007). We calculated the rhabdom acceptance angle Δ*ρ* (Fig. 4B) according to (Land et al., 1997),

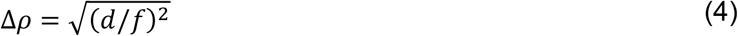

The sensitivity N of each ommatidium (Fig. 4A) was calculated according to (Warrant & Nilsson, 1998) as proportional with the following eye parameters:

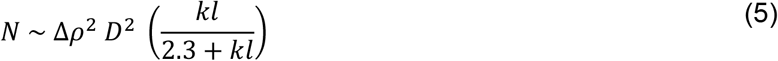

We calculated the eye parameter *P* (Fig. 4F) as a measure for the investment of the eye in sensitivity (larger values) or acuity (smaller values) according to (Snyder, 1977):

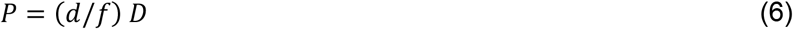

Where *λ* is the light’s wavelength, set to 500 nm, *k* is the absorption coefficient of the photoreceptor (0.0067 μm^-1^), *D* is the facet diameter, *d* is the rhabdom diameter, *l* is the rhabdom length, and *f* is the focal length of the eye. The focal length of the eye cannot be determined directly from anatomical measures in aspherical superposition compound eyes (Warrant, 1999). We could assume the same scaling coefficients for the focal length as for the eye diameter for the following reason: the focal length in superposition compound eyes is principally measured as the distance from the nodal point of the eye to the tip of the retina (Land et al., 1997). The nodal point of the eye depends on the radius of the eye, which scaled isometrically with eye size (Fig. 1H, S3F). The distance from the nodal point to the tip of the retina depends on the radius of the eye, and the distance of the retina to the cornea. The latter remained constant across animals of different sizes (Fig. S5), so that the scaling of the focal length was determined by the eye radius, which in turn scaled isometrically with eye diameter. We therefore used the eye diameter scaling coefficient for the focal length and fitted the intercept to an average focal length of 0.375 mm (Warrant et al., 1999).

We used the computational estimations of eye function to assess how the descriptors of eye function varied with different allometric scaling. To this aim, we calculated the ommatidial sensitivity (equation 5) and rhabdom acceptance angle (equation 4), as well as the eye parameter (equation 6) for a range of allometric scaling exponents across many eye sizes (corresponding to the body sizes in Fig. 4A-C). Since the focal and rhabdom lengths scaled isometrically, and the facet and rhabdom diameters scaled with very similar exponents (Figs. 2D, 3C), we calculated the eye performance measures by applying the same scaling exponent factor to both the facet and rhabdom diameter, while retaining the focal and rhabdom lengths at isometry (Fig. 4D-F). We then assessed the variance of the given eye function descriptors across eye size for each scaling exponent as the standard deviation (Fig. 4G-I).

## Acknowledgements

The authors would like to thank Pierre Tichit for support with data acquisition and discussion of data analysis, and Ola Gustafsson, Eva Landgren and Carina Rasmussen for support with the sample preparation. We would like to thank Eric Warrant for helpful discussion of the optical modelling. We acknowledge funding to A.S. by the Würzburg Graduate School of Life Sciences PostdocPlus Programme and the German Research Council (DFG: STO 1255 2-1), and to E.B. by the Air Force Office of Scientific Research (FA8655-12-1-2136), the Vetenskapsrådet (2014-4762) and the Carl Tryggers Stiftelse för Vetenskaplig Forskning (CTS15:38).

## Supplementary Figures

**Fig. S1.**
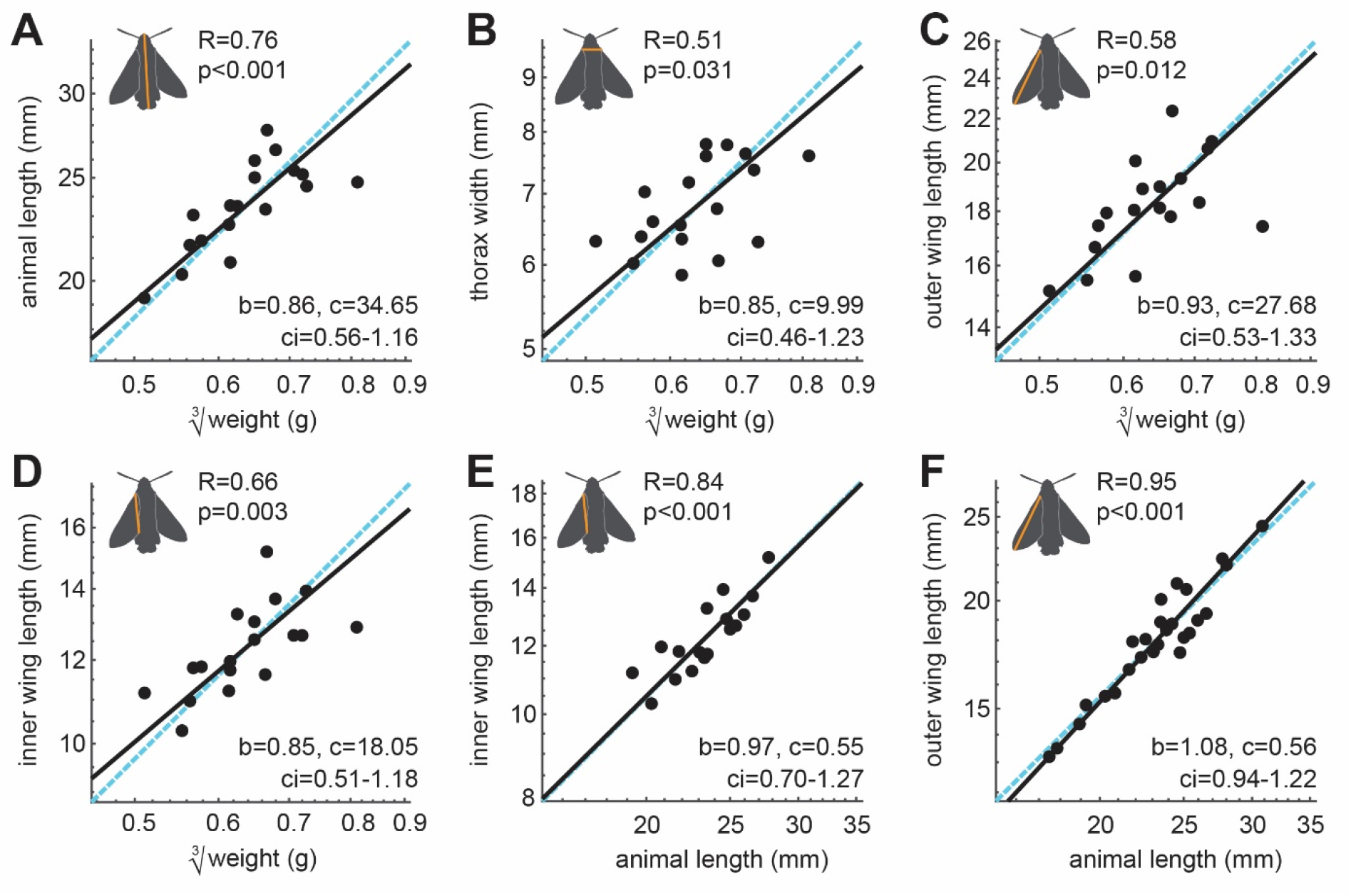
Allometric scaling of body size in *Macroglossum stellatarum*. Allometric scaling of the **A** anterior-posterior body length, **B** the thorax width, **C** the outer wing length, **D** and the inner wing length with the cube-root of body weight. **E** Scaling of the inner and **F** outer wing length with animal length. Measurements are indicated by the orange lines. **A**-**F** Data from individual hawkmoths (black dots). The dashed cyan line indicates isometric scaling and the black line represents the allometric scaling relationship. *R* is the Pearson correlation coefficient of the log-transformed data, and *p* denotes its statistical significance. Given the significant linear correlations in **A**-**F**, the allometric relationship was calculated using reduced major axis regression, with the exponential scaling exponent *b*, the normalization constant *c*, and the confidence interval *ci* of *b*.

**Fig. S2.**
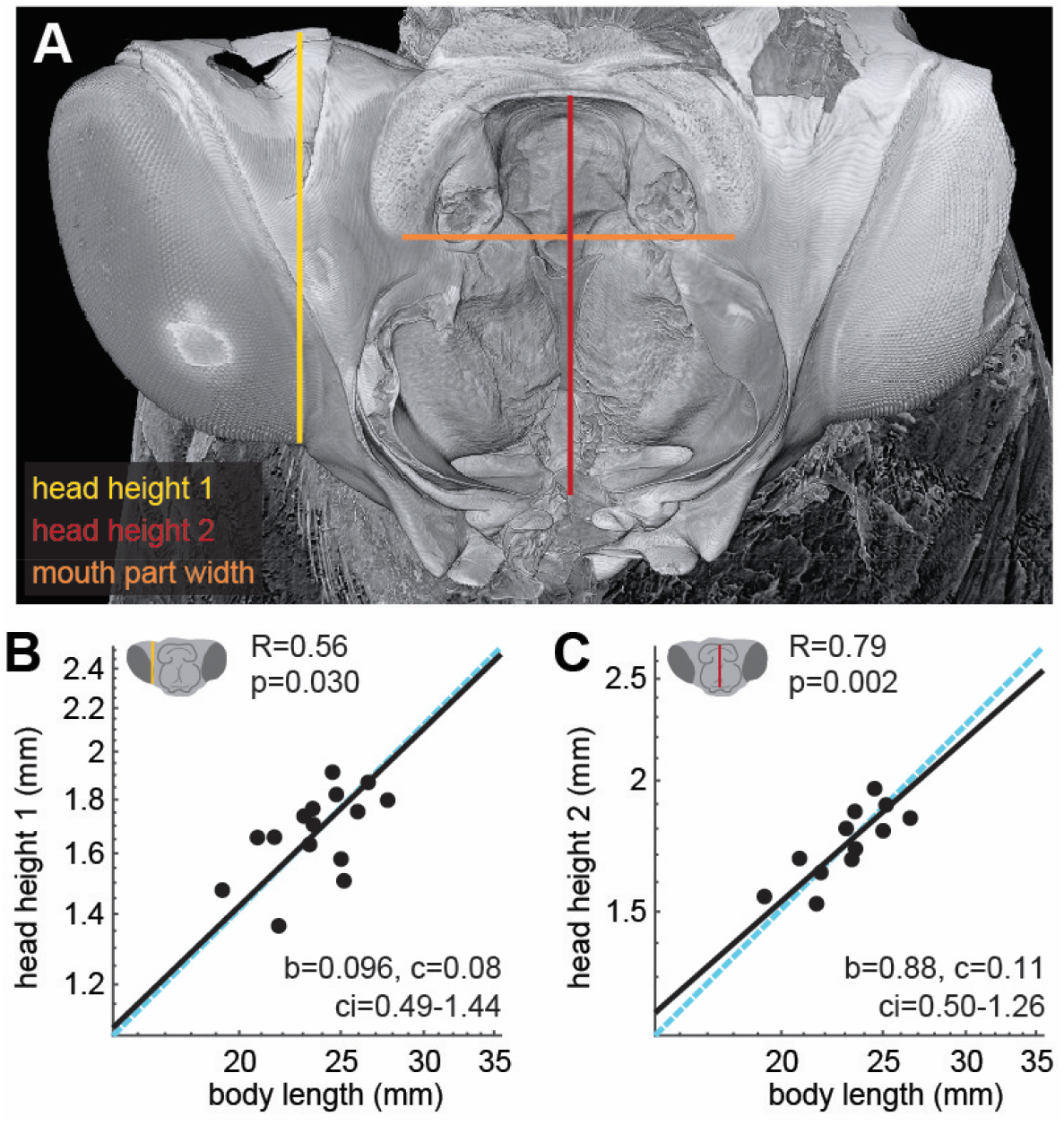
Allometric scaling of head size with body size. **A** Microtomography images of the head of *M. stellatarum*. The left eye was cut open for better penetration of the fixative, and the mouth parts, dorsal and posterior face of the head were removed as well. To assess head size, we measured the lateral extend of the mouth part base (1, Fig. 1G), the dorso-ventral extent of the mouth part base (2) and the dorso-ventral extent of the right optic lobe (3). Allometric scaling of the **B** optic lobe height, and **C** the height of the mouth-part base with body length. **B**-**C** Data from individual hawkmoths was measured by X-ray microtomography (black dots). The dashed cyan line indicates isometric scaling and the black line represents the allometric scaling relationship. *R* is the Pearson correlation coefficient of the log-transformed data, and *p* denotes its statistical significance. Given the significant linear correlations in **A**-**F**, the allometric relationship was calculated using reduced major axis regression, with the exponential scaling exponent *b*, the normalization constant *c*, and the confidence interval *ci* of *b*.

**Fig. S3.**
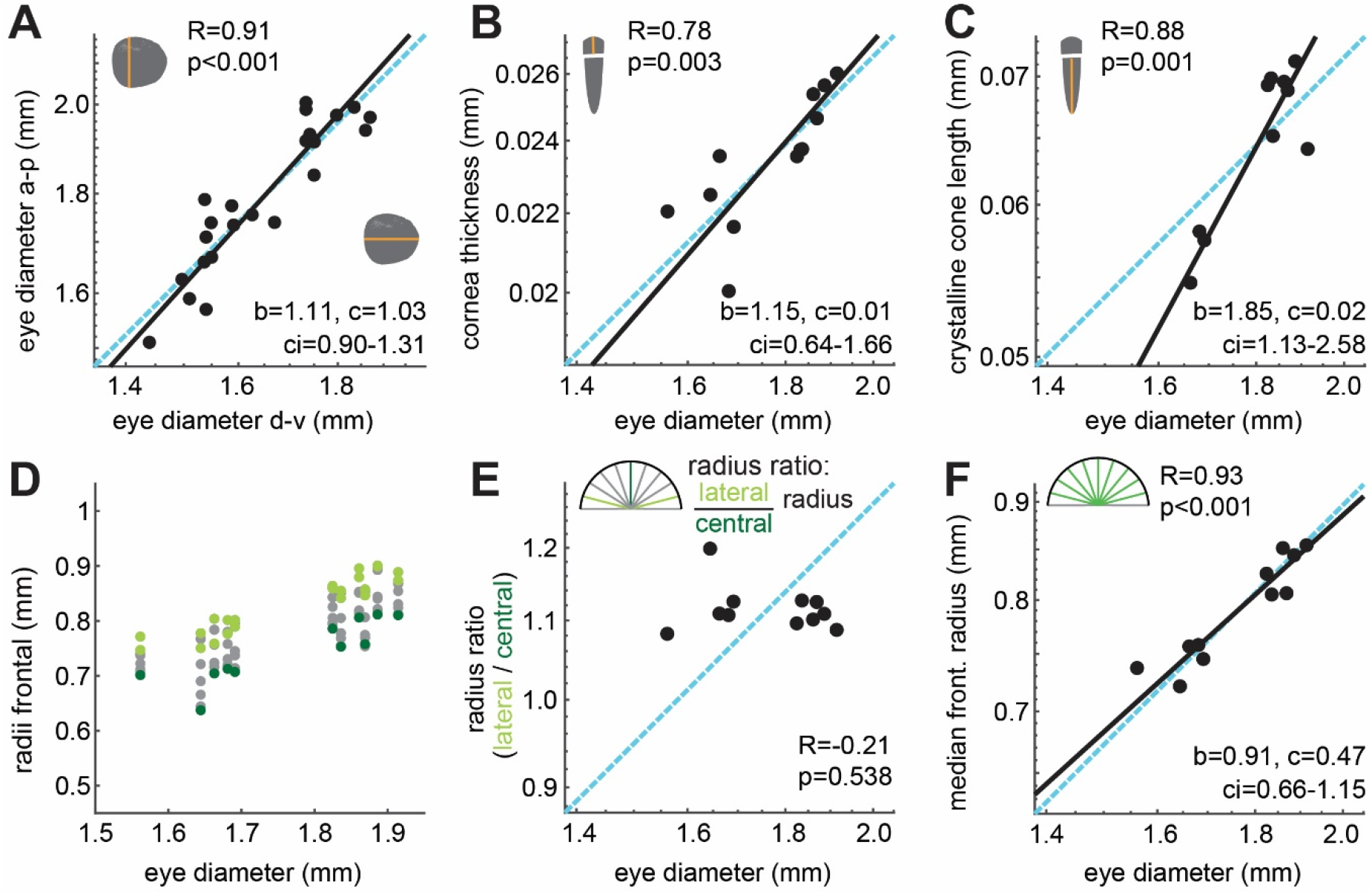
Allometric scaling cornea structures and corneal shape with eye size. Allometric scaling of the **A** anterior-posterior with dorso-ventral eye diameter, and **B** of the facet thickness **C** crystalline cone length with eye size. **D** To test whether the curvature of the eye differed across eye diameters, we measured the distance from the nodal point formed by the edges of the cornea to the corneal surface for nine evenly spaced radii in frontal sections. **E** We calculated the ratio between the average lateral radii (light blue) and the central radius (dark blue) as a proxy for the corneas’ shape and assessed its allometric scaling. **F** shows the allometric scaling of the median of the central seven radius measurements (green) with eye diameter. **A**-**C**,**E**,**F** Data from individual hawkmoths (black dots). The dashed cyan line indicates isometric scaling and the black line represents the allometric scaling relationship. *R* is the Pearson correlation coefficient of the log-transformed data, and *p* denotes its statistical significance. Given the significant linear correlations in **A**-**C**,**F**, the allometric relationship was calculated using reduced major axis regression, with the exponential scaling exponent *b*, the normalization constant *c*, and the confidence interval *ci* of *b*.

**Fig. S4.**
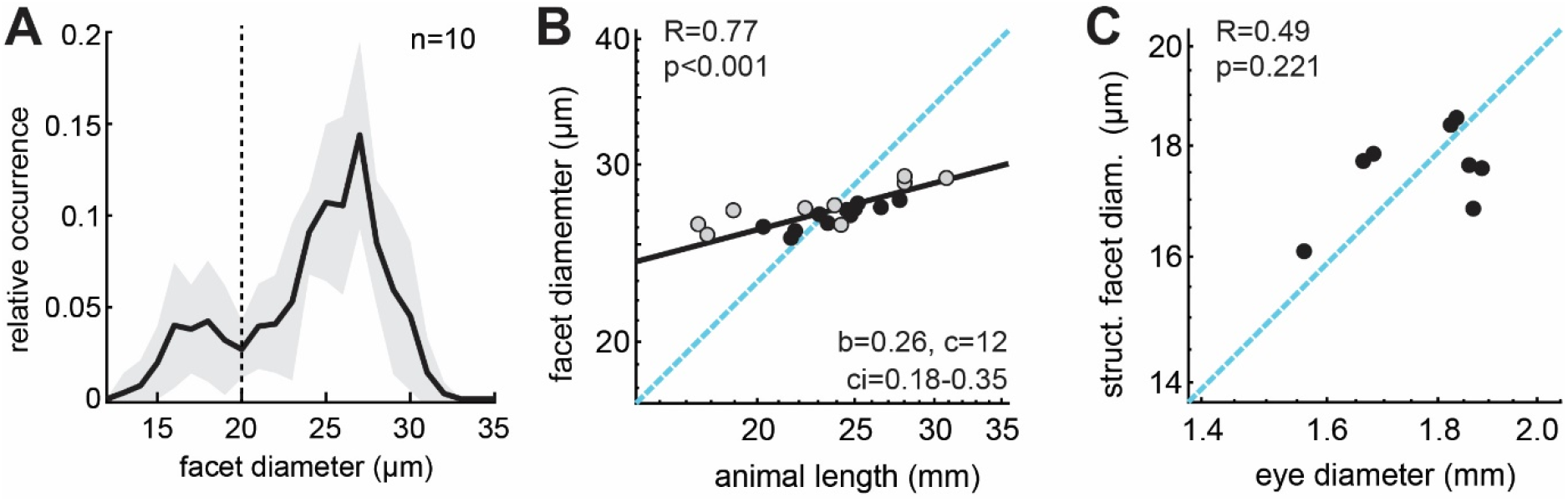
Allometric scaling of structural and functional facets. **A** Histograms of all facet diameters measured across the corneal surface of 10 eyes. The black line represents the mean and the shaded area the standard deviation. The dashed line indicates the threshold for separating structural from functional facets at 20 μm. Allometric scaling **B** of the functional facet diameter with animal length, and **C** the structural facet diameter with eye diameter. **B**-**C** Data from individual hawkmoths was measured by either X-ray microtomography (black dots), or light-microscopy (grey dots). The dashed cyan line indicates isometric scaling and the black line represents the allometric scaling relationship. *R* is the Pearson correlation coefficient of the log-transformed data, and *p* denotes its statistical significance. Given the significant linear correlations in **B**, the allometric relationship was calculated using reduced major axis regression, with the exponential scaling exponent *b*, the normalization constant *c*, and the confidence interval *ci* of *b*.

**Fig. S5.**
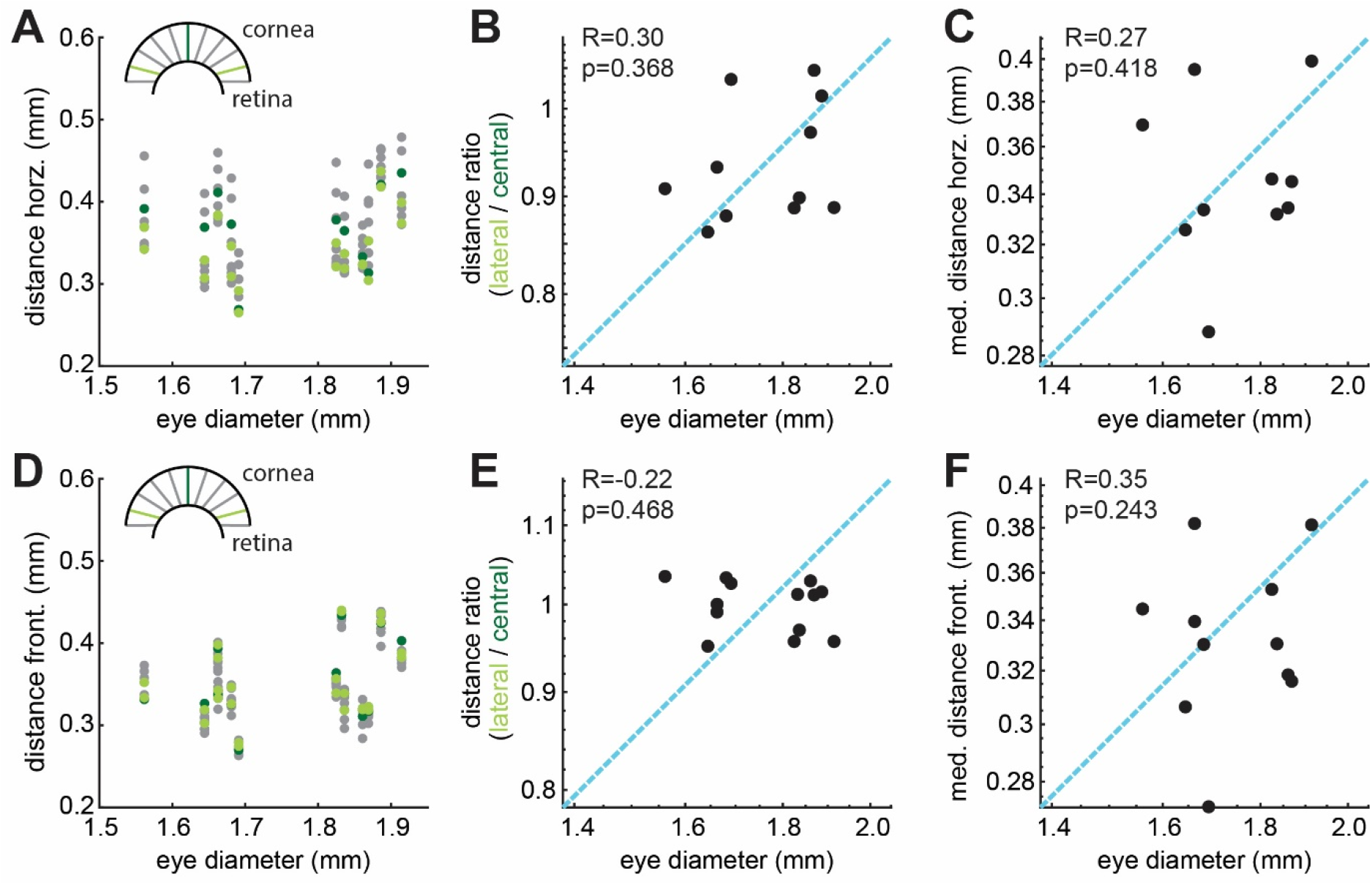
Allometric scaling of retina-cornea distance. To test whether the distance of the cornea to retina differed across eye diameters, we measured this distance for eleven evenly spaced measurements in **A** horizontal and **D** frontal sections. Allometric scaling of the ratio of lateral and central measurements with eye diameter from **B** horizontal and **E** frontal sections, and of the median **C** horizontally and **F** frontally measured distance between retina and cornea with eye diameter. **B**-**C**,**E**-**F** Data from individual hawkmoths was measured by X-ray microtomography (black dots). The dashed cyan line indicates isometric scaling and the black line represents the allometric scaling relationship. *R* is the Pearson correlation coefficient of the log-transformed data, and *p* denotes its statistical significance.

**Fig. S6.**
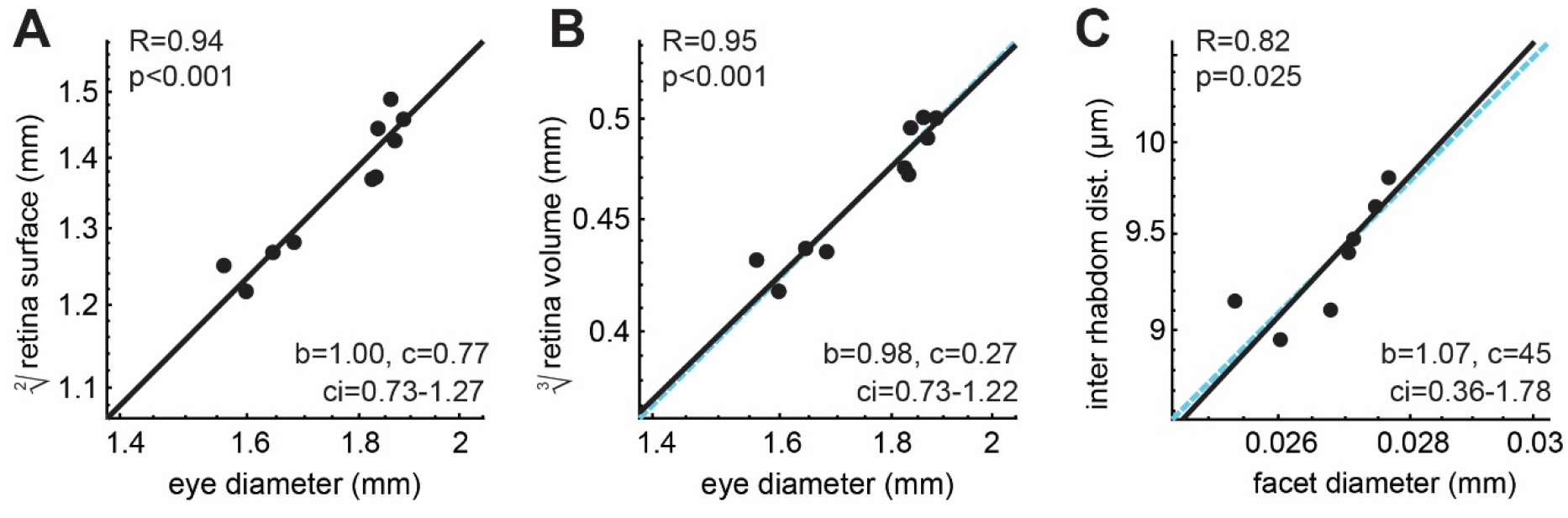
Allometric scaling of the retinal surface, volume and rhabdom distance. Allometric scaling **A** of the square root of the retina surface area obtained by 3D reconstructions, **B** the cube-root of retinal volume with eye size, and **C** and the relationship between of the rhabdom distance with and facet diameter. **A**-**C** Data from individual hawkmoths was measured by X-ray microtomography (black dots). The dashed cyan line indicates isometric scaling and the black line represents the allometric scaling relationship. *R* is the Pearson correlation coefficient of the log-transformed data, and *p* denotes its statistical significance. Given the significant linear correlations in **B**, the allometric relationship was calculated using reduced major axis regression, with the exponential scaling exponent *b*, the normalization constant *c*, and the confidence interval *ci* of *b*.

